# Circular RNA profiling reveals abundant and diverse circRNAs of SARS-CoV-2, SARS-CoV and MERS-CoV origin

**DOI:** 10.1101/2020.12.07.415422

**Authors:** Shaomin Yang, Hong Zhou, Ruth Cruz-Cosme, Mingde Liu, Jiayu Xu, Xiaoyu Niu, Yaolan Li, Lizu Xiao, Qiuhong Wang, Hua Zhu, Qiyi Tang

## Abstract

Circular RNAs (circRNAs) encoded by DNA genomes have been identified across host and pathogen species as parts of the transcriptome. Accumulating evidences indicate that circRNAs play critical roles in autoimmune diseases and viral pathogenesis. Here we report that RNA viruses of the *Betacoronavirus* genus of *Coronaviridae*, SARS-CoV-2, SARS-CoV and MERS-CoV, encode a novel type of circRNAs. Through *de novo* circRNA analyses of publicly available coronavirus-infection related deep RNA-Sequencing data, we identified 351, 224 and 2,764 circRNAs derived from SARS-CoV-2, SARS-CoV and MERS-CoV, respectively, and characterized two major back-splice events shared by these viruses. Coronavirus-derived circRNAs are more abundant and longer compared to host genome-derived circRNAs. Using a systematic strategy to amplify and identify back-splice junction sequences, we experimentally identified over 100 viral circRNAs from SARS-CoV-2 infected Vero E6 cells. This collection of circRNAs provided the first line of evidence for the abundance and diversity of coronavirus-derived circRNAs and suggested possible mechanisms driving circRNA biogenesis from RNA genomes. Our findings highlight circRNAs as an important component of the coronavirus transcriptome.

**Summary:** We report for the first time that abundant and diverse circRNAs are generated by SARS-CoV-2, SARS-CoV and MERS-CoV and represent a novel type of circRNAs that differ from circRNAs encoded by DNA genomes.

## INTRODUCTION

Severe acute respiratory syndrome coronavirus 2 (SARS-CoV-2) is a single strand and positive sense RNA virus and belongs to the *Betacoronavirus* genus of the family of *Coronaviridae* (CoVs). It is responsible for the ongoing global pandemic of COVID-19. SARS-CoV-2 shares ~80% homology with severe acute respiratory syndrome coronavirus (SARS-CoV) and is more closely related with Middle East respiratory syndrome-related coronavirus (MERS-CoV) than other four commonly circulated human coronaviruses (*1, 2*). SARS-CoV-2, SARS-CoV and MERS-CoV, emerged within last two decades and have posed major challenges to global health. However, we still have very limited understanding of their pathogenicity factors. The transcriptional regulation of CoV gene expression is complex due to the large size of the genome (~30kb). The first open reading frame (ORF), ORF1a/1b, is translated from the positive-strand genomic RNA (gRNA) as a polyprotein, which is cleaved proteolytically into non-structural proteins. ORFs located towards the 3’ side of the genome encode conserved structural proteins, including S (spike protein), E (envelope protein), M (membrane protein) and N (nucleocapsid protein), and accessory proteins. These proteins are translated from a set of sub-genomic RNAs (sgRNA) generated through TRS-L and TRS-B (transcription-regulating sequences from the leader and body) mediated discontinuous RNA synthesis (*3*). It is recently revealed that the transcriptome of SARS-CoV-2 is even more complex with numerous non-canonical discontinuous transcripts produced and potentially encoding unknown ORFs through fusion, deletion, truncation and/or frameshift of existing ORFs (*4*). It is unclear if additional components exist in the transcriptome of SARS-CoV-2 and other CoVs.

Circular RNAs (circRNAs) are a class of single-stranded noncoding RNA species with a covalent closed circular configuration. CircRNAs are formed either through back-splicing of exons or from intron lariat by escaping debranching (5). CircRNAs are resistant to exonuclease-mediated degradation and are more stable than linear RNA (*6*). They may encode proteins (*7*) or function as miRNA and protein sponges (*8*). Recent studies have revealed circRNAs as important pathological biomarkers for cancers (*9*), neurological diseases (*10*) and autoimmune diseases (*11*). Furthermore, viral-derived circRNAs have been identified from several DNA viruses, including Epstein-Barr Virus (*12–14*), Kaposi Sarcoma Virus (*15–17*) and human papillomaviruses (*18*), and are implicated with a role in pathogenesis (*18*).

In this study, we report the bioinformatical identification and characterization of SARS-CoV-2-, SARS-CoV- and MERS-CoV-derived circRNAs as a novel type of circRNAs using publicly available deep RNA-Seq data. We also present the first systematic approach to validation circRNAs expressed by SARS-CoV-2. We experimentally identified over 100 circRNAs, which supports the major findings from our bioinformatic analyses. Our results demonstrate the abundance and diversity of circRNAs derived from RNA viral genomes of beta-coronaviruses, providing insights into the biogenesis and functions of circRNAs during viral infection.

## RESULTS

### Identification of SARS-CoV-2-, SARS-CoV- and MERS-CoV-derived circRNAs and characterization of back-splice junction hotspots using CIRI2

It is recommended that bioinformatic analyses of circRNAs are performed on datasets with at least 30 million 100-bp raw reads generated from cDNA libraries prepared from rRNA-depleted total RNA (*19*). To look for circRNAs derived from CoV genomes, we identified SARS-CoV-2-, SARS-CoV- and MERS-CoV-infection-related deep RNA-Seq datasets in the NCBI Gene Expression Omnibus database. Considering the replication kinetics and tropism of CoVs (*20*), we chose datasets from GSE153940 (*21*), GSE56193, and GSE139516 (*22*), with 24 hours post infection (hpi) as the timepoint, Vero E6 (African green monkey kidney) cells as the host for SARS-CoV-2 and SARS-CoV, and Calu-3 (human lung adenocarcinoma) cells as the host for MERS-CoV. A circRNA enrichment step was included during cDNA preparation for the MERS-CoV datasets (*22*), rendering the MERS-CoV datasets more sensitive for circRNA detection.

CoVs use an RNA-dependent RNA polymerase (RdRp) to generate genomic RNA and sgRNA transcripts in the cytoplasm of host cells. We thus reasoned that CoV circRNAs, if existed, are likely to circularize independent of splicing, which occurs in the nucleus. Several circRNA prediction algorithms have been developed to identify BSJ reads from RNA-Seq data and to predict the 5’ and 3’ breakpoints (*23*). CIRI2 (*23*) is the only tool that adopts an MLE-based algorithm to unbiasedly identify back-splice junction (BSJ) reads independent of a circRNA reference annotation file. It is more sensitive and accurate than two other *de novo* circRNA identification tools (*23*). Therefore, we used the recommended CIRI2 pipeline (*24*) to perform *de novo* circRNA discovery and assembly.

To improve the assembly accuracy and to simplify follow-up comparison, we combined reads of biological triplicates into single datasets. After mapping with BWA-MEM (*25*), we obtained 1,216,403,242 total reads from the SARS-CoV-2 dataset with 36.6% mapped to SARS-CoV-2. The MERS-CoV dataset had a similar percentage (30.2% of 316,893,928 total reads) mapped to the viral genome. And 87.0% of the 1,127,121,362 total reads from the SARS-CoV dataset was mapped to SARS-CoV. The SARS-CoV-2 and SARS-CoV datasets showed sharp peaks at the 5’ leader sequence and high coverage towards the 3’ end of the genome (Figure 1A and 1B). Genome coverage of the MERS-CoV dataset was substantially lower due to the removal of linear RNAs by RNase R (Figure1A and 1B). We observed above-threshold coverage in the last 5,000 nucleotides (nt) of the MERS-CoV genome, corresponding to E, N, ORF8b and the 3’UTR.

**Figure 1.**
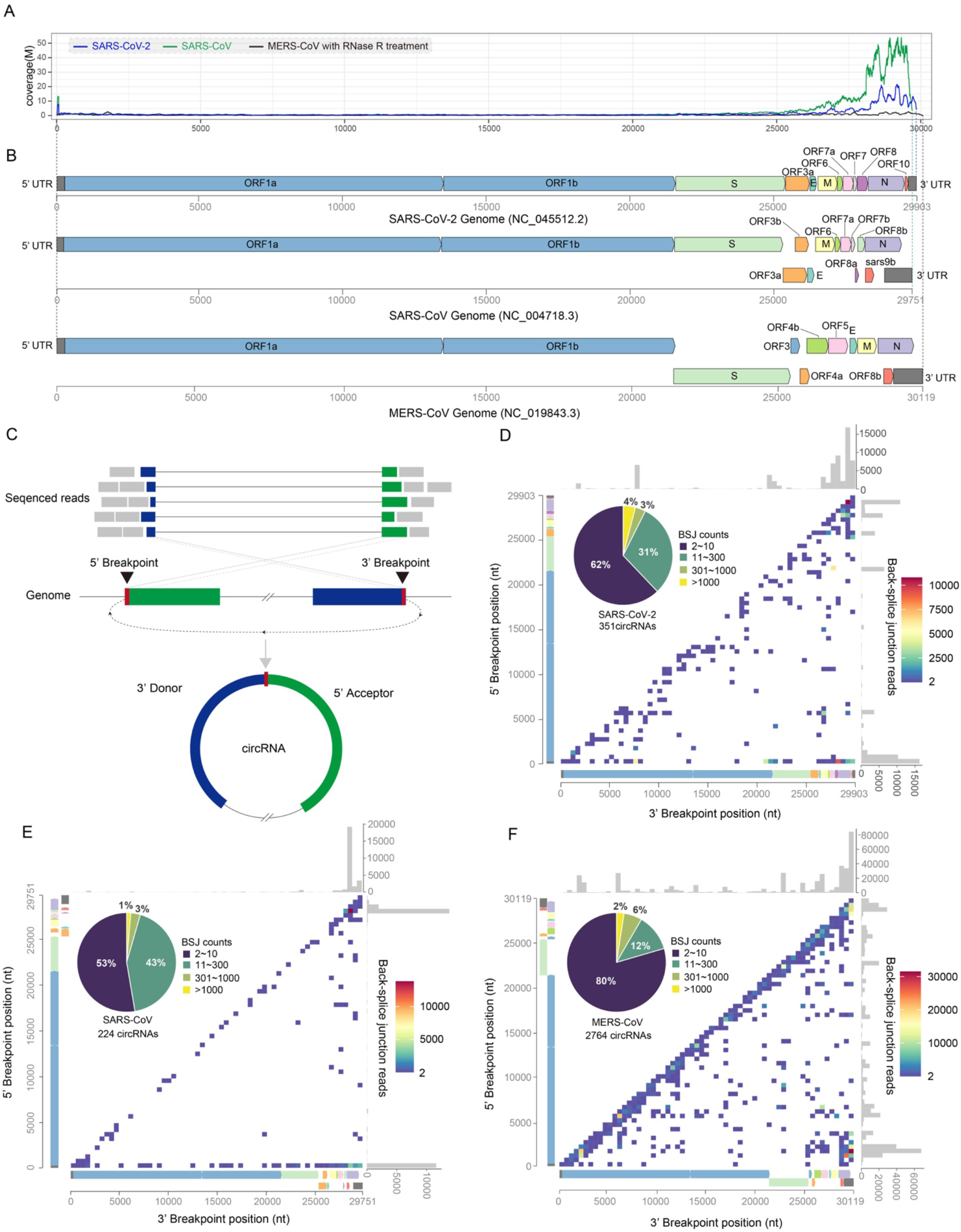
Identification of SARS-CoV-2-, SARS-CoV- and MERS-CoV-derived circRNAs. **(A)** Coverage of SARS-CoV-2, SARS-CoV and MERS-CoV genomes in CoV-infected related deep RNA-Seq data. **(B)** Genome organization of SARS-CoV-2, SARS-CoV and MERS-CoV. **(C)** Illustration of BSJ-spanning reads aligned to the donor and acceptor sequences, and determination of the 5’ and 3’ breakpoints. The relative locations of breakpoints in the linear and circular RNAs are shown. **(D-F)** Frequency of circularization events in SARS-CoV-2 (D), SARS-CoV € and MERS-CoV (F). Counts of BSJ-spanning reads (starting from a coordinate in the X axis and ending in a coordinate in the y axis) indicated by color. The counts were aggregated into 500nt bins for both axes. Distribution of start/end position was shown as histograms on the x and y axis. The number of identified circRNAs from each CoV genome and the breakdown of read counts was shown as pie charts.

CIRI2 identifies circRNAs by aligning chimeric reads to the 3’ donor sequence and the 5’ acceptor sequence and determining the exact breakpoints of the BSJ (Figure 1C). By this definition, we identified 351 SARS-CoV-2 circRNAs, 224 SARS-CoV circRNAs and 2,764 MERS-CoV circRNAs. The larger number of circRNAs identified from MERS-CoV genome compared to SARS-CoV2 and SARS-CoV demonstrates the efficiency of circRNA enrichment with RNase R digestion. While the majority of CoV-derived circRNAs had very low (<10) BSJ-spanning reads, 14 SARS-CoV-2 circRNAs (4%), 3 SARS-CoV circRNAs (1%) and 68 MERS-CoV circRNAs (2%) had over 1,000 BSJ-spanning reads (Figure 1D-1F and S1F). An additional 3-6% of the identified circRNAs had 300-1,000 BSJ-spanning reads (Figure 1D-1F). In fact, the most abundant circRNA identified in each CoV dataset had >10,000 BSJ-spanning reads (SARS-CoV-2_29122|29262: 10,763; SARS-CoV_28136|28606: 13,690; MERS-CoV_1503|29952: 29,467). While more circRNAs were identified from the host genomes (monkey: 10,291; human: 43357), the overall expression level of host circRNAs is much lower compared to CoV circRNAs (Figure S1F).

To examine the circRNA landscape, we mapped all identified circRNAs by the 5’ and 3’ breakpoints of the BSJs to their respective genomic locations and estimated the back-splicing frequency by counting the reads spanning the BSJs (Figure 1D-1F). We identified two major types of back-splicing events shared by all three CoVs: 1) long-distance back-splicing between the 3’ end of the genome and the 5’ end of the genomes; 2) local back-splicing in regions corresponding to the N gene of SARS-CoV-2 and SARS-CoV and the 3’UTR of MERS-CoV). We also noticed back-splicing events that specifically occur in SARS-CoV-2 or MERS-CoV. Local back-splicing around position 1500-2500 (Nsp2), 5500-6500nt (Nsp3) and 22000-23000nt (S) of the MERS-CoV genome occurred at high frequency (Figure 1F), whereas middle-distance back-splicing from SARS-CoV-2 genomic region 7501-8000 (Nsp3) to 1-500 (5’UTR) and from 27501-28000 (ORF7a/ORF7b) to 22001-22500nt (S) was observed at high frequency (Figure 1D).

Next, we performed *de novo* reconstruction and quantification of full-length SARS-CoV-2, SARS-CoV and MERS-CoV circRNAs using the CIRI-full (*24*) algorithm. We got 300 reconstructed SARS-CoV-2 circRNAs, of which 127 (42.3%) were full-length. Of 201 assembled SARS-CoV circRNAs, 122 (60.7%) were full-length. We also got 1,024 reconstructed MERS-CoV circRNAs, with 81.6% were fully assembled, suggesting that RNase R treatment improves circRNA reconstruction. *De novo* assembly of host circRNAs resulted in 4,815 (49.9%) full-length monkey circRNAs and 31,808 (100%) full-length human circRNAs.

Furthermore, we compared the features of circRNAs derived from CoVs with those from the host genomes. The length of nuclear genome-derived circRNAs (nu-circRNAs) is highly conserved across species with the majority ranging from 250 to 500 nt (*24*). We observed similar length distribution in full-length monkey and human genome-derived circRNAs (Figure 2A). CoV circRNAs shared a different length distribution pattern (Figure 2B). The average length of SARS-CoV-2 and MERS-CoV circRNAs was over 150 nt longer than that of the host circRNAs (Figure 2A and 2B). And more SARS-CoV-2 and MERS-CoV circRNAs were over 1,000 nt long whereas host circRNAs are rarely over 750 nt in length. Since CoV have both positive and negative genomic and subgenomic RNAs, we examined the strandness of CoV circRNAs. CircRNAs generated by both host genomes showed no strand preference (Vero: 51.9% positive-stranded; Calu-3: 51.0% positive-stranded). In contrast, 59.5% of SARS-CoV-2 circRNAs, 56.3% of SARS-CoV circRNAs, and 85.1% of MERS-CoV circRNAs were negative-stranded (Figure 2A). This result suggests that CoV circRNAs have a preference for negative strand.

**Figure 2.**
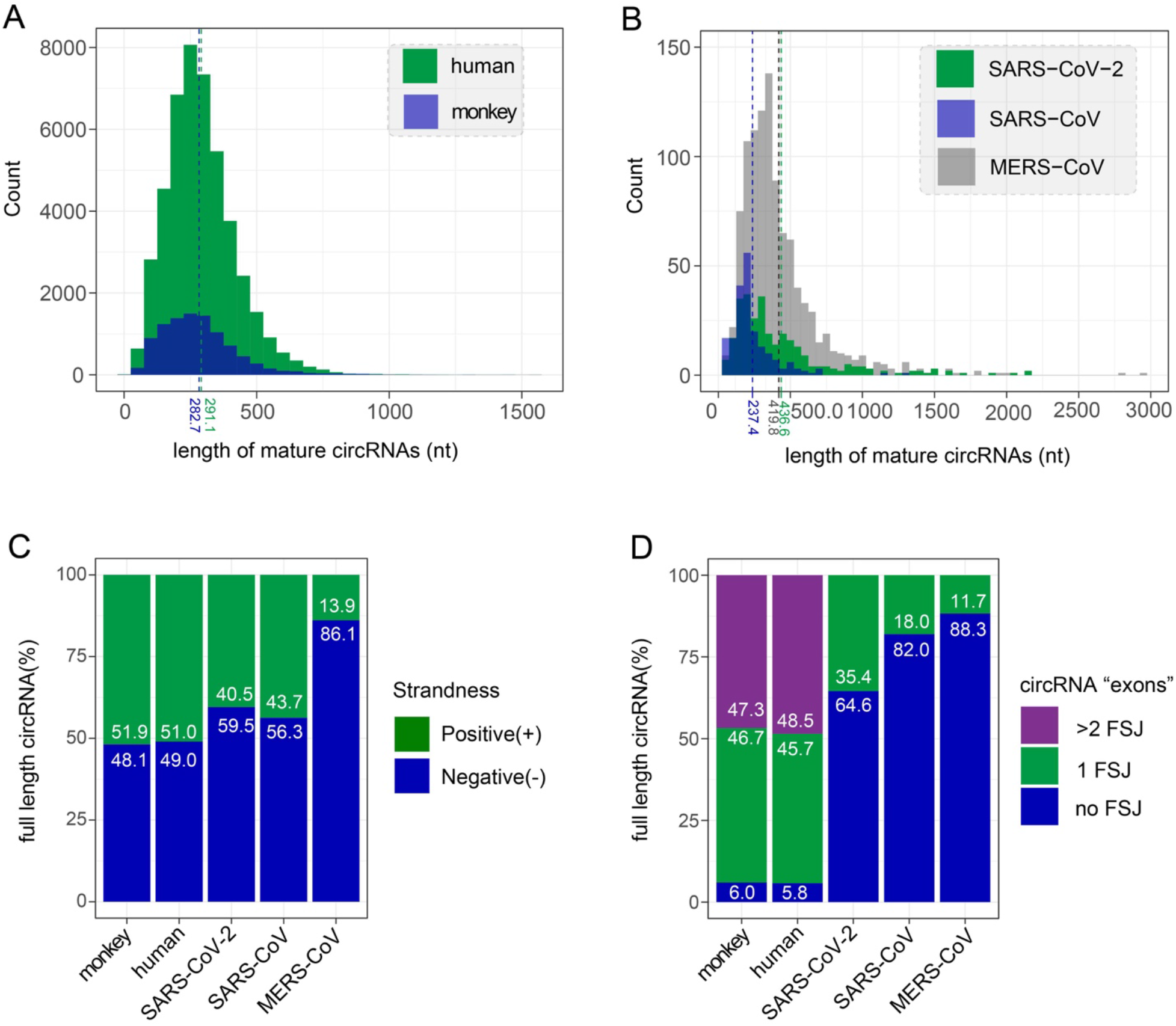
Comparison of predicted full-length CoV circRNAs and host circRNAs. **(A) and (B)** Length distribution of circRNAs derived from host genomes (A) and CoVs (B). Average length indicated by dashed lines. **(C)** Strand distribution of host and viral circRNAs. **(D)** Distribution of circRNA exons in host and viral circRNAs. Only full-length circRNAs predicted by CIRI2-full were quantified.

Nu-circRNAs with the same BSJ often have a diverse number of forward-splicing junctions (FSJs) and circRNA exons due to alternative intron retention (*24*). sgRNA with canonical and non-canonical FSJs have been observed in CoVs (*3, 4*), suggesting that CoV circRNAs may also have FSJs and circRNA isoforms. We examined the number of FSJs in full-length host and CoV circRNAs. While circRNA without FSJ only represent 6% of host circRNAs, the majority of CoV circRNAs had no FSJ (SARS-CoV-2: 64.6%; SARS-CoV: 82%; MERS-CoV: 83.3%). Additionally, only 1 FSJ could be detected in predicted full-length CoV circRNAs, whereas about 50% of host circRNAs had at least 2 FSJs (Figure 2D). Next, we looked for predicted full-length CoV circRNAs that share the same BSJ breakpoints but differ in length. We found that MERS-CoV circRNA 1262|29148 produces two isoforms, both of which contain one FSJ. The longer isoform (1,051nt) has the FSJ 2223|29060, whereas the shorter isoform (155nt) has the FSJ 1316|29049. This result shows that very few CoV circRNAs could have isoforms.

In conclusion, we analyzed SARS-CoV-2, SARS-CoV and MERS-CoV related deep RNA-Seq datasets, and identified a large amount of CoV circRNAs. The circRNAs of CoV origin have features in common and can be distinguished from circRNAs derived from the human and monkey host genomes. We have shown that CoV circRNAs are expressed at higher level and longer in length than host circRNAs and tends to be negative stranded. We identified BSJ hotspots for circRNAs derived from each CoV, and found that distant back-splicing from the tail of the genome to the head of the genome and local back-splicing in regions corresponding to the N gene and the 3’UTR occur at the highest frequency.

### Experimental detection and analysis of SARS-CoV-2 circRNAs

We extracted total RNA from Vero E6 cells mock-treated or infected with SARS-CoV-2 at 24 hpi. Forward and reverse divergent primers were designed to maximize the chances of amplifying BSJ sequences (Figure 3A and 3B). To validate the two major back-splicing events, we performed inverse RT-PCR with primer pairs that targeting either the distant BSJ hotspot 29001-29903|1~500 or the local BSJ hotspots 28501~29500|27501~28500 (Figure S2A-S2C). We also performed inverse RT-PCR with divergent primer sets targeting the most abundant SARS-CoV-2 circRNAs predicted by CIRI2 (Figure 3C). Majority of the inverse RT-PCR reactions using the infected sample as template resulted in products ranging from 200bp to 800bp, whereas no amplification was seen from the mock samples. Notably, many candidate inverse RT-PCR products were more abundant than that of circHIPK3, a known highly expressed human circRNA that served as a positive control (Figure 3C, S2A and S2B). We gel-purified candidate PCR products based on the size, subcloned by TA cloning, and Sanger-sequenced at least 8 colonies for each candidate BSJ sequence. The sequencing results revealed the surprising diversity of SARS-CoV-2 circRNAs and support our predictions from the bioinformatic analyses. First, all gel-purified bands represent more than one PCR product of the same size. While highly expressed circRNAs, such as 29194|27797 and 28853|28467, represent over 50% of the confirmed clones (29194|27797: 5/7 with 29083-F and 27893-R; 28853|28467: 4/8 with 28809-F and 28494-R; Figure 3D and 3F), most other purified bands contain a variety of circRNAs (data not shown). Secondly, we confirmed that the breakpoints of a given circRNA is surprisingly flexible. For example, PCR products amplified by 29668-F/29572-F and 51-R contain a distant BSJ. However, the 3’ breakpoint ranges from genomic location 29,080nt to 29,767nt, and the 5’ breakpoint was between genomic location 7nt and 19nt (Figure S3B). When a deviation of 10nt was considered for the breakpoints, the predicted BSJ 29758|8 represent 8 out of the 13 BSJs confirmed by sequencing. Thirdly, both the distant and the local back-splicing events were validated by multiple BSJs. We detected distant fusion from ORF6, N, ORF10 and the 3’UTR to the 5’ UTR (data not shown). We also detected local fusion within N, and from N to ORF7a, ORF7b, and ORF8 (data not shown). In summary, our RT-PCR and sequencing results validated the diversity of SARS2 circRNAs at the genome level and at the circRNA level.

**Figure 3.**
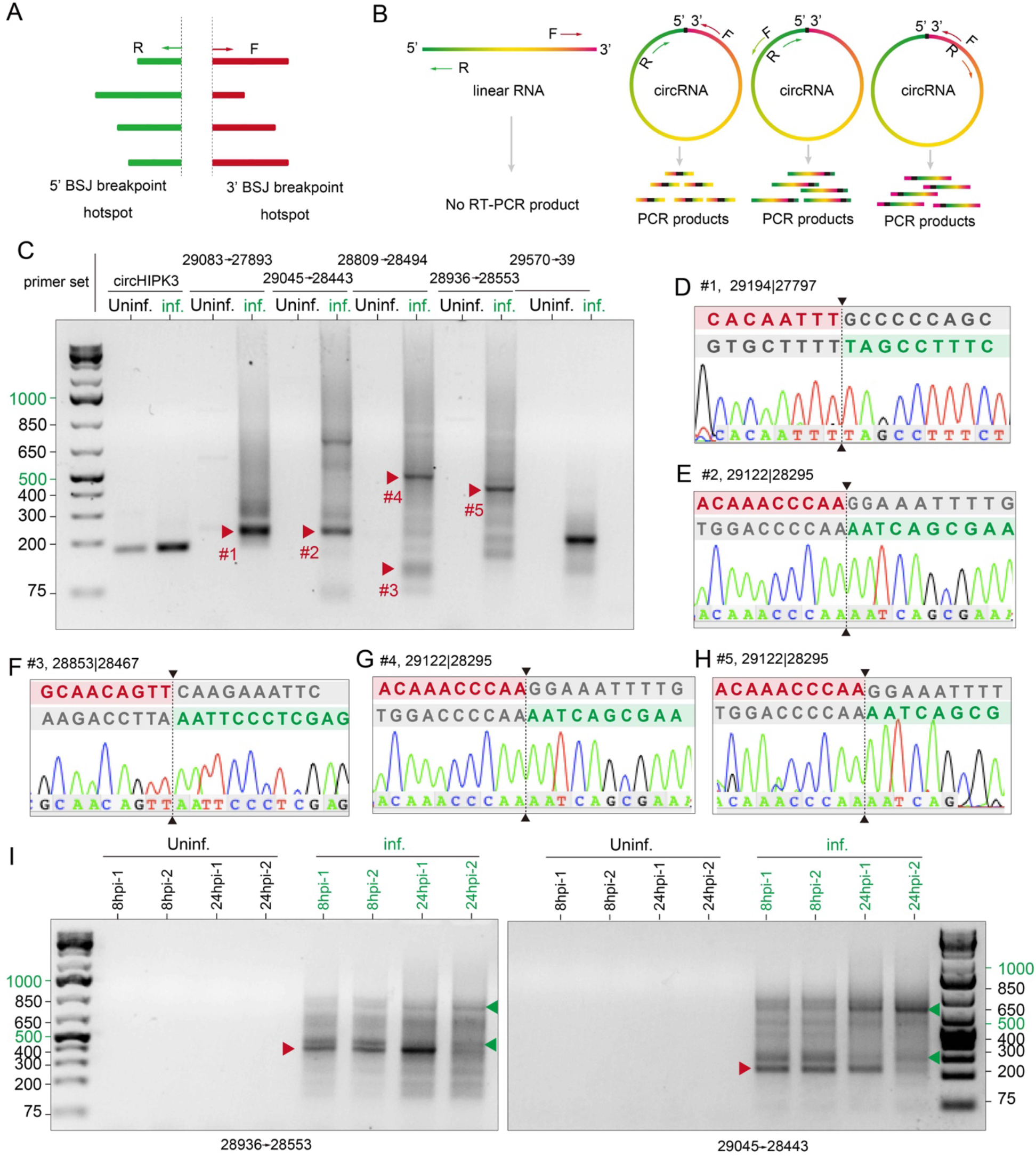
Experimental validation of SARS-CoV-2 circRNAs in Vero E6 cells. **(A)** Schematic showing divergent primers were designed to amplify all predicted BSJs in a given hotspot. **(B)** Illustration of BSJ RT-PCR with divergent primers would selectively amplify different regions of circRNAs but not linear RNAs. **(C)** BSJ RT-PCR with selected primer sets. Bands indicated by red arrows were gel-purified and sequenced. Note the intensity of most candidate BSJs were comparable to that of the positive control, circHIPK3 of host origin. Infection also enhanced the expression of circHIPK3. **(D-H)** Examples of Sanger sequencing results for PCR products in (C). Sequences around the 3’ and 5’ breakpoints were aligned to the BSJ sequence. BSJ Breakpoints were indicated by dashed lines. Donor and acceptor sequences were highlighted in magenta and green, respectively. Sequences excluded from the circRNA were shown in grey. **(I)** BSJ RT-PCR probing SARS-CoV-2_29122|28925 in uninfected and infected Vero E6 cells at 8hpi and 24hpi. Primer sets were labelled at the bottom of the gels. Red arrows correspond to bands #5 and #2 in (C). Green arrows indicate candidate circRNAs that are differentially expressed at early and late stage of infection.

While the inverse RT-PCR was designed to amplify sequences around the BSJs, we successfully assembled the full-length sequence of circRNA 29122|28295, of 828nt in length, using a combination of primer sets (29045-F/28443-R, 28486-F/28341-R, 28809-F/28494-R and 28642-F/28553-R). The successful detection of circRNA 29122|28295 with multiple primer pairs (Figure 3C, 3E, 3G and 3F) and the high rate of detection in subclones (data not shown) indicate the overwhelming abundance of this circRNA. In fact, this circRNA corresponds to the most abundant SARS-CoV-2 circRNA 29122|29262 predicted by CIRI2. This result demonstrates the accuracy of our bioinformatic analysis.

To better understand the consistency of SARS-CoV-2 circRNA expression, we probed SARS-CoV_29122|28295 in biological replicates of uninfected and infected samples at 8hpi and 24hpi with two divergent primer sets. RT-PCR with a convergent primer pair targeting the N gene confirmed that the viral titer was comparable among the infected samples (Figure S3A). We found that the bands (red arrowheads) corresponding to circRNA 29122|28295 were strong in all the samples except for infected-24hpi-rep2, which is still detectable but significantly lower (Figure 3I). Interestingly, we found that the abundance of others candidate BSJ products (green arrows) amplified by these primer sets was different between 8hpi and 24hpi samples. This result suggests that circRNA expression level and pattern could change over the course of infection.

We also confirmed a few features of CoV circRNAs characterized bioinformatically. First, we detected a variety of FSJs in SARS-CoV-2 circRNAs. The major type of FSJ was accompanied with a long-distance back-splicing to the 5’UTR to create sgRNA-like circRNAs. We found 5 circRNAs that contained FSJ 75|28266 and 4 circRNAs that contained FSJ 76|26480 (data not shown), suggesting TRS-mediated fusion of the leader sequence with N and M gene, respectively. Interestingly, the BSJs in sgRNA-like circRNAs were more flexible. The 3’ breakpoints ranges from 28465 to 2927, and the 5’ breakpoint ranges from 3 to 40 (Figure S3B). It is likely that these circRNAs used sgRNAs as template for synthesis. We also detected FSJs that represent noncanonical “splicing” events. 6066|29068 and 15466|28579 are long-range TRS-L-independent distant fusion, whereas 28353|28408, 28353|28471, and 28666|28729 represent noncanonical local fusions in the N genes, all of which are consistent with recent finding of noncanonical fusion in the SARS-CoV-2 transcriptome (*4*). Secondly, we confirmed alternative back-splicing events in SARS-CoV2 circRNAs either with shared 5’ breakpoints or shared 3’ breakpoints. Distant back-splicing from various loci in the N gene share the same 5’ breakpoints in the 5’UTR, such as 28465|40 and 29273|40. Fusion from the 3’ end of the M gene (genomic location 27282nt) to either the TRS-L (47nt) or TRS-B (26484,) was observed.

Two circRNAs with unexpected repetitive back-splicing caught our attention. One had two different distant back-splicing events (28465|40 and 28526|1) followed by the same TRS-L dependent fusion, 75|28266 (Figure S3C). The other had two rounds of fusion from 28465 to 28320 followed by a third fusion from 28467 to 28282 (Figure S3D). Since the BSJs within the same circRNAs were slightly different, it is unlikely to be an artifact of the rolling-cycle amplification of circRNAs by RT. These two cases suggest that SARS-CoV-2 circRNAs form BSJs independent of splicing. It is likely that SARS-CoV-2 circRNA are generated through the template-switching mechanism that drives the formation of discontinuous transcripts. In support of this hypothesis, we found that the upstream sequences of the acceptors were homologous to the donor sequence (Figure 3D-H, data not shown). TRS-dependent FSJs in SARS-CoV-2 circRNAs had 11-12 homologous nucleotides between the leader and the body sequence. Also, BSJs with 3-6 nucleotides homology around the breakpoint was frequently observed.

In conclusion, we have demonstrated that SARS-CoV-2 produces a surprising diversity of circRNAs that are abundantly present in the infected Vero E6 cells.

## DISCUSSION

CircRNAs are a recently discovered and recognized type of RNA with important roles in diseases. While some studies have been conducted in the context of viral infection, the focus was on how host circRNAs respond to infection. So far, only limited viral circRNAs have been identified from viruses, mostly from large DNA viruses of the family of *herpesviridae*, and the circular RNA genome of the hepatitis delta virus is the only known closed circRNAs produced by an RNA virus (*26*). Here we provide the first line of evidence that RNA genomes of beta-coronaviruses encode a novel type of circRNAs, which differ from those encoded by DNA genomes. In this study, we took two approaches: 1) bioinformatically profiling of the circRNA landscape in SARS-CoV-2, SARS-CoV and MERS-CoV as well as their human and African green monkey hosts by *de novo* circRNA identification and assembly of public available deep RNA-Seq datasets using CIRI2; 2) experimentally profiling of the circRNA landscape in SARS-CoV-2 by systematic capturing and identifying viral circRNAs produced from the predicted BSJ hotspots.

We bioinformatically identified 351, 224 and 2,764 circRNAs derived from SARS-CoV-2, SARS-CoV and MERS-CoV, respectively (Figure 1D-1F), and experimentally identified more than 100 SARS-CoV-2 circRNAs (data not shown). Comparing the BSJ landscapes and frequency among SARS-CoV-2, SARS-CoV and MERS-CoV revealed two major circularization events shared by all the three CoVs: 1) distant fusion between RNA located at the tail and the head of the genome; 2) local fusion in the conserved N gene (Figure 1D-1F). These events were confirmed by experimentally identified circRNAs (Figure 3C-H and S3B). What distinguishes CoV circRNAs from host circRNAs are the expression level (Figure S1F), the length (Figure 2A and 2B), the strand preference (Figure 2C), and the circRNA exon number (Figure 2D).

The collection of experimentally identified SARS-CoV-2 circRNAs further distinguishes CoV circRNAs from Nu-circRNAs. First, we observed striking flexibility in the breakpoints of SARS-CoV-2 circRNAs. Analysis of sequences around the 3’ and 5’ breakpoints of experimentally identified SARS-CoV circRNAs suggest that homology-mediated inaccurate fusion drives the back-splicing event (data not shown), whereas nu-circRNAs tend to splice accurately on the AGGT splicing signal. Secondly, we found two cases where multiple back-splicing events occurred in the same circRNAs (Figure S3C and S3D), suggesting back-splicing occurs as the RNA is synthesized. It further suggests that the RNA configuration could create BSJ hotspots that enable repetitive back-splicing.

As we wrote this manuscript, another group reported the first bioinformatic identification of circRNAs in SARS-CoV-2, SARS-CoV and MERS-CoV (*27*). Interestingly, they came to several opposing conclusions about CoV circRNAs, including the abundance, the strandness and the expression level. It is likely due to the datasets they used and the circRNA analysis pipeline and strategy they adopted. First, we chose SARS-CoV-2 and SARS-CoV datasets with higher sequencing depth and pooled biological triplicates before the analysis. As a result, we identified 240 circRNAs shared by CIRI2 and finc_circ (Figure S1E), twice the number they found. Since CoV circRNA does not form BSJs through splicing, AG|GT signal-base algorithms are likely to have an extreme high false discovery rate, which could lead to their opposing conclusion on strand-preference. Secondly, we chose BSJ-spanning read counts as the indication of abundance and made comparison between the host and the viral circRNAs of the same dataset. We have shown that many CoV circRNAs were spliced tail-to-head. Using transcript per million (TPM) as the index would greatly underestimate the abundance of CoV circRNAs. Similarly, they considered the span between the 5’ and 3’ breakpoints of the BSJ is the length of the circRNA, assuming that CoV circRNAs do not have FSJs, is an unreasonable way to analyze the data. For our analysis, we only quantified fully assembled circRNAs predicted by CIRI2-full, rendering our length analysis more reliable. Lastly, the group claimed that the number of circRNA identified by their pipeline increased over the course of infection. However, our experimental results suggest that the most abundant SARS-CoV-2 circRNA, 29122|28295, was highly expressed at 8 hpi and was likely to down-regulated at 24 hpi (Figure 3I). Considering the flexibility of circRNA BSJs, we have observed experimentally and the inaccuracy of bioinformatic algorithms in calling circRNAs. We believe using a systematic approach to examine circRNA expression diversity and abundance at different stages of infection is needed before any conclusion could be drawn.

Taken together, we have demonstrated with bioinformatic analyses and experimental evidence that a novel class of circRNAs are generated from SARS-CoV-2, SARS-CoV and MERS-CoV genomes. The CoV circRNA are highly diverse and abundant, comprising an important part of the CoV transcriptome. Our study provide insight into the biogenesis of CoV circRNA and the functions of CoV circRNAs during pathogenesis and viral replication. Understanding the nature and biological function of CoV circRNAs will help us to understand how these viruses evade the host immune system, replicate and course diseases.

## AUTHOR CONTRIBUTIONS

S.Y and H.Z. designed the experiments, S.Y, H.Z., R.C., M.L., J.X., X.N., Q.T., performed the experiments, S.Y., H.Z., H.Z., Q.T, analyzed the data, H.Z., H.Z., Q.T., Q.W. wrote the paper, Y.L., L.X, Q.W, H.Z., Q.T, supervised the study.

## ACKNOWLEDGEMENTS

This study was supported by an NIH/NIAID SC1AI112785 (Q.T.), an NIH/DE R01DE028583-01 (subaward to Q.T.), and National Institute on Minority Health and Health Disparities of the National Institutes of Health under Award Number G12MD007597.

The following reagent was deposited by the Centers for Disease Control and Prevention and obtained through BEI Resources, NIAID, NIH: SARS-Related Coronavirus 2, Isolate USA-WA1/2020, NR-52281. We thank Dr. Juliette Hanson and Kaitlynn Starr for BSL3 training and assistance in BSL3-related work. Q.W. and her group were supported by state and federal funds appropriated to Ohio Agricultural Research and Development Center (OARDC), College of Food, Agricultural, & Environmental Sciences, The Ohio State University.

## METHODS AND MATERIALS

### *De novo* circRNA identification and reconstruction

The analysis workflow was performed on two Intel W-3175X CPUs with 128 GB memory running Ubuntu system (version 18.04)(*28*). Adaptor trimmed reads of the same condition were pooled and aligned with BWA Aligner(*25*) (BWA-MEM version 0.7.17-R1188) and bowtie2 (version 2.3.5.1)(*29*) to host and viral reference genomes: Afircan green monkey (ChlSab1.1.101) for bioproject PRJNA168621; human (hg19) for bioproject PRJNA31257; SARS-CoV-2 (NC_045512.2) for bioproject PRJNA485481; SARS-CoV (NC_004718.3) for bioproject PRJNA485481; and MERS-CoV (NC_019843.3) for bioproject PRJNA485481. Alignment statistics was performed with Qualimap2 (version 2.2.1)(*30*). CIRI2 (version v2.0.6)(*23*) and find_circ (version 1.2) (*31*) were used for circRNA calling. Reconstruction of partial and full length circRNAs was performed with CIRI-full (version 2.0)(*24*). Default setting was used.

### Quantification and plotting

Quantification and plots were produced using python (version 3.9.0) with plotly module (https://plotly.com/python/ and R statistical environment (version 3.4.5) with R package: gggenes (https://wilkox.org/gggenes/, Figure 1B), ggplot2 (other Figures)(*32*).

### Cell culture, plasmid DNA transfection and SARS-CoV-2 infection

Vero cells (ATCC, CCL-81) and HEK 293T(ATCC® CRL-1573™) were purchased from ATCC. The cells were maintained in Dulbecco’s modified Eagle’s medium (DMEM) supplemented with 10% fetal calf serum (FCS) and penicillin (100 IU/ml)-streptomycin (100 ug/ml) and amphotericin B (2.5 ug/ml) (*33*).

The plasmid, pCAG-nCoV-N-FLAG (*34*) expresses nucleocapsid (N) gene and was transfected into HEK 293T cells by transfection reagent, Lipofectamine 3000 (cat# L3000015, Scientific Fisher, USA) according to the manufacturer’s protocol.

The SARS-CoV-2 infection experiment was performed in BSL3 labs as described previously (*35*). Eight T75 flasks of Vero E6 cells (ATCC No. CRL-1586) formed 90-100% confluency were used. After washing with DMEM (Life Technologies) twice, four flasks of cell monolayers were inoculated with SARS-CoV-2 USA-WA1/2020 strain (BEI Resources, NIAID, NIH), which has been passaged one time in Vero E6 cells after we received it from BEI Resources, diluted in 15 mL of DMEM supplemented with 2% of heat inactivated (56°C for 30min) fetal bovine serum (Hyclone) and 100 units penicillin/mL, 100 μg streptomycin/mL, and 0.25 μg amphotericin B/mL (Sigma). We used a multiplicity of infection (MOI) of 0.3 based on 50% tissue culture infectious dose (TCID50). The other four flasks were incubated with medium only as mock. At 8 hours post-inoculation (hpi) and 24 hpi, we stopped incubating half of the virus-inoculated and mock flasks by gently pipetting out the culture supernatant. Then we added 5 mL TRIzol™ (Invitrogen) into each flask and gently rocked the flasks to distribute the Trizol solution evenly. After pipetting several times to remove all cells, we transferred the lysates to chloroform-resistance tubes. After keeping the tubes in room temperature for 5 min to fully lysis the cells, we took 100 μL/sample for inactivation test by performing two rounds of virus isolation in Vero E6 cells. The rest of the samples were stored at −80°C. After the validation of virus inactivation, the samples were moved out of BSL3 facility for circRNA analyses in BSL2 laboratories.

### Experimental detection and analysis of SARS-CoV-2 circRNAs

Detection and analysis of SARS-CoV-2 circRNAs was performed as previously described (*36*). Total RNA was isolated using TRizol (ThermoFisher) and Direct-zol RNA miniprep kit (Zymo) from mock-treated and SARS-CoV-2-infected Vero E6 cells at 8hpi and 24 hpi. RNase R (Lecigen) treatment and follow-up purification (RNA Clean and Concentrator, Zymo) was performed as described in (*36*). If RNase R treatment is opted out, 500ng total RNA was used for reverse transcription (Superscript IV, ThermoFisher) with random hexamer primers (ThermoFisher). Divergent and convergent primers used in this study are summarized in Table S1. PCR was performed with GoTaq Master Mix (Promega) with 1ul cDNA template at 1:20 dilution. Following agarose gel (2%) electrophoresis, candidate circRNA PCR products were size-selected and gel-purified (Gel purification kit, Zymo) and subcloned with TA cloning kit (ThermoFisher). At least 8 colonies were checked for insertion of candidate PCR products by PCR with M13 universal primers. Amplified insertions were PCR purified (DNA purification kit, Zymo) and subjected to Sanger sequencing by MCLAB, CA. Sequencing results were blasted against SARS-CoV-2 reference genome (NC_045512.2). 5’ and 3’ breakpoints of BSJs and FSJs were manually curated. All commercial reagents were used according to manufacturer instruction.

