## supplemental data for "Circular RNA profiling reveals abundant and diverse circRNAs of SARS-CoV-2, SARS-CoV and MERS-CoV origin"

**Fig. S1**

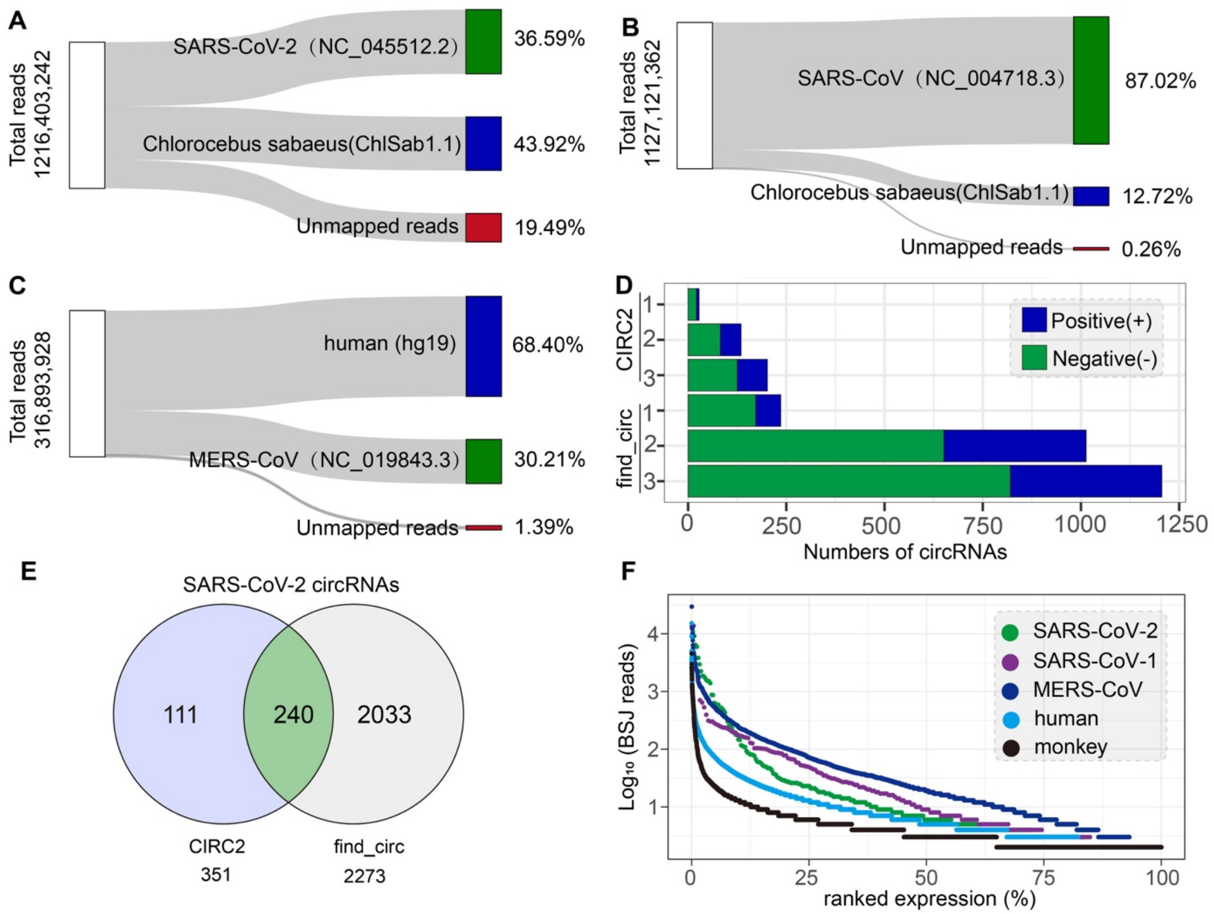


**Figure S1. Related to Fig. 1. (A-C)** Statistics of pooled RNA-Seq data. Total reads and mapping efficiency to host and viral genomes as shown. **(D)** Comparison of circRNAs identified by CIRI2 and find_circ from the SARS-CoV-2 datasets. RNA-Seq data of biological replicates were analyzed separately. **(E)** Number of circRNAs identified by both CIRI2 and find_circ or only by one algorithm. **(F)** Ranked expression level of all *de novo* identified CoV and host circRNAs.

**Fig. S2**

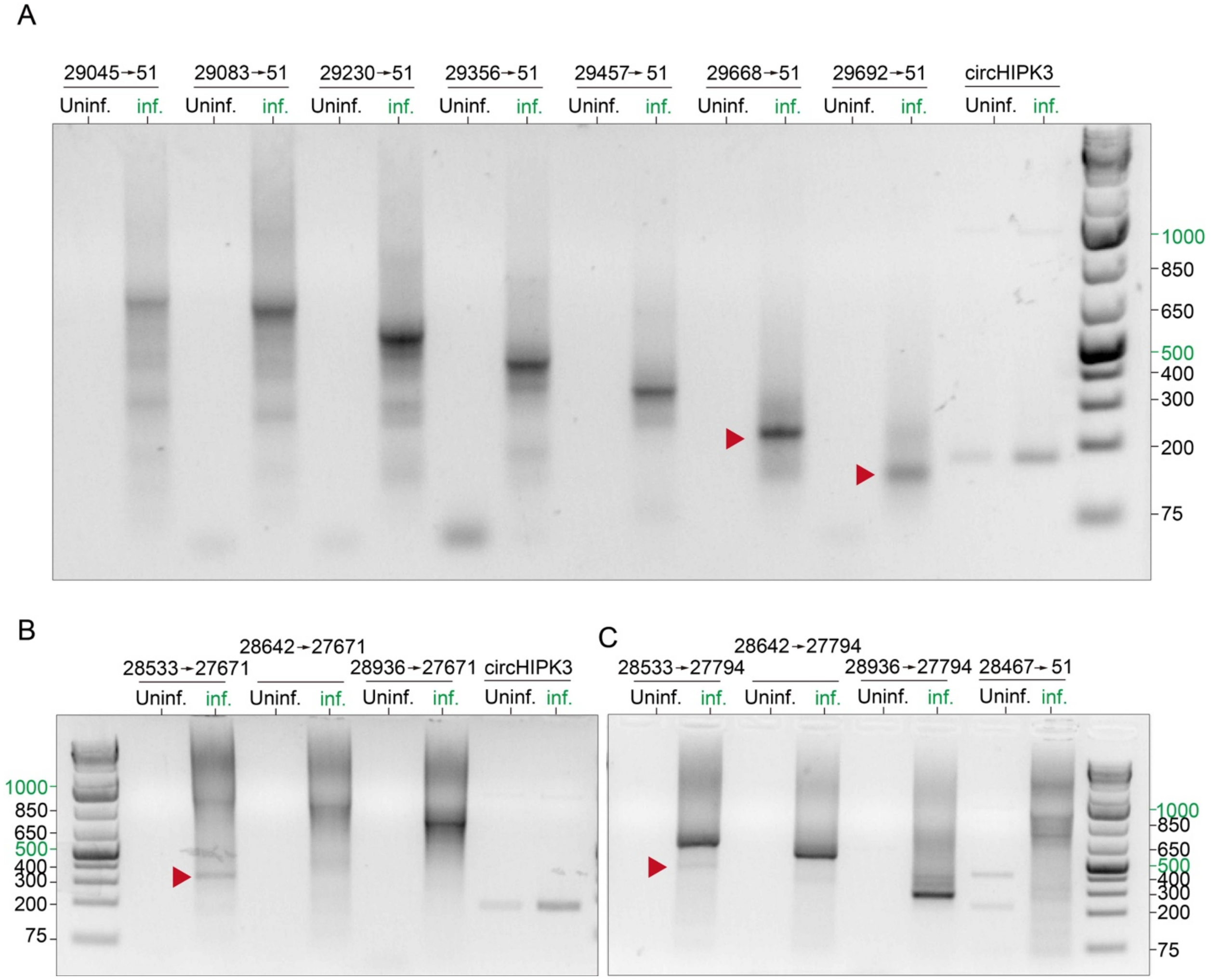


**Figure S2. Systematic amplification of candidate SARS-CoV-2 circRNAs.** Primer sets as indicated. Candidate circRNA in the distant BSJ hotspot 29001-29903|1~500 **(A)** or the local BSJ hotspots 28501~29500|27501~28500 **(B-C)** were shown. Green arrows indicate Sanger sequencing identified circRNAs in **Figure S3B**.

**Fig. S3**

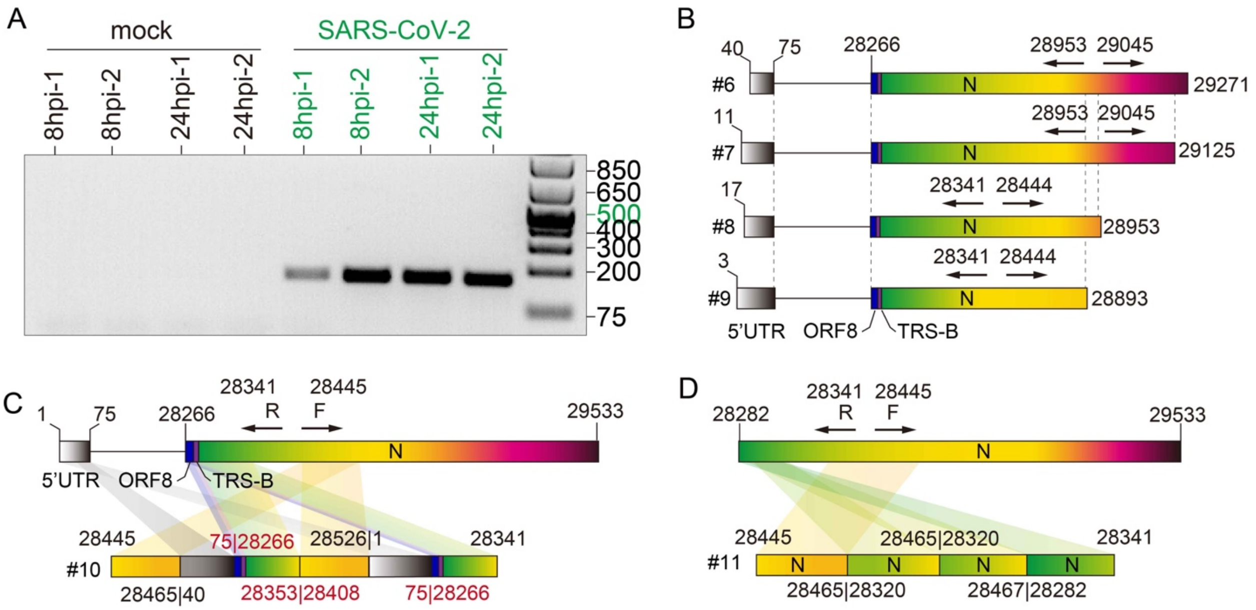


**Figure S3. Related to Fig. 3 and Fig. S2. (A)** Validation of SARS-CoV-2 infection efficiency. Convergent primers against N was used. Refer to Table S1 for details. **(B)** Illustration of sequencing-validated circRNAs from **Fig. S2**. Primers as indicated. **(C)** and **(D)** represent two cases of confirmed circRNAs with more than one BSJs.

**Table S1. Primers used in this study**

| Name | Sequence (5' to 3') |
|---|---|
| **Divergent primers** | |
| 39-R | TTGGTTGGTTTGTTACCTGGG |
| 51-R | AGAGATCGAAAGTTGGTTGGT |
| 27671-R | ACTTCCTCTTGTCTGATGAACA |
| 27794-F | GCCTTTCTGCTATTCCTTGTTT |
| 27893-R | GTTCGTTTAGGCGTGACAAGT |
| 28443-R | GTGAGAGCGGTGAACCAAGA |
| 28467-F | CAACATGGCAAGGAAGACCT |
| 28494-R | ATTGGAACGCCTTGTCCTCG |
| 28533-F | ACCGAAGAGCTACCAGACGA |
| 28553-R | TTCGTCTGGTAGCTCTTCGGT |
| 28642-F | TGGTGCTAACAAAGACGGCAT |
| 28809-F | GCAGTCAAGCCTCTTCTCGT |
| 28936-F | GCTGCTGCTTGACAGATTGA |
| 29045-F | CCTCGGCAAAAACGTACTGC |
| 29083-F | AACACAAGCTTTCGGCAGAC |
| 29230-F | CATTGGCATGGAAGTCACAC |
| 29356-F | AACATTCCCACCAACAGAGC |
| 29457-F | TTCCTGCTGCAGATTTGGAT |
| 29570-F | AACGTTTTCGCTTTTCCGTTT |
| 29668-F | CACATAGCAATCTTTAATCAGTGTG |
| 29692-R | CACACTGATTAAAGATTGCTATGTGA |
| circHIPK3-F | TTCAACATATCTACAATCTCGGT |
| circHIPK3-R | ACCATTCACATAGGTCCGT |
| **Convergent primers** | |
| N-F(29083-F) | AACACAAGCTTTCGGCAGAC |
| N-R (29249-R) | GTGTGACTTCCATGCCAATG |
